# The effect of external lateral stabilization on energy cost of walking – A systematic review and meta-analysis

**DOI:** 10.1101/2021.09.07.459220

**Authors:** Mohammadreza Mahaki, Marco J M Hoozemans, Han Houdijk, Jaap H. van Dieën, Sjoerd M Bruijn

## Abstract

Previous studies have estimated the energy cost required for the control of medio-lateral stability in human walking by means of external lateral stabilization. Results were inconsistent, possibly due to differences in task constraints or stabilization devices. To better understand the effects of lateral stabilization on energy cost, we conducted a systematic review with meta-analysis of studies, which directly assessed effects of lateral stabilization on energy cost in healthy young adult participants (18-41 years old). We obtained individual participant data on net energy cost (J kg^-1^ m^-1^) from previously published studies. Across all studies reviewed, the net energy cost reduction during stabilized walking at preferred and zero step widths equaled to 0.05 ± 0.35 (~2-3% reduction) and 0.25 ± 0.29 J kg^-1^ m^-1^ (mean ± s.d.) (~8-9% reduction), respectively. The effect of external lateral stabilization was significant only for walking at zero step width and without arm swing. Lateral stabilization devices with short rope length increased energy cost reduction. However, spring stiffness and habituation time did not influence energy cost reduction. We provide recommendations for improvement of lateral stabilization devices to avoid some of the confounding effects. External lateral stabilization reduces energy cost during walking by a small amount. It can be concluded that a small proportion of total energy cost is required to control medio-lateral stability; this proportion is larger when walking with narrow steps and without arm swing.

## 1. Introduction

Humans walk in ways that minimize energy cost [1–3]. For instance, humans prefer to walk at a speed or to select a step frequency which makes walking less energetically costly [1, 3]. It has been shown that aging [4] or neurological disorders [5, 6] may increase this energy cost which subsequently may make walking a strenuous task and limit the mobility. Hence, it is important to understand what constitutes the costs of human walking.

During walking, humans consume metabolic energy to generate propulsive force [7], to initiate and propagate leg swing [8] and to maintain stability [9, 10]. Gottschall and Kram estimated the energy cost of propulsion at nearly 50% [7] and the cost of generating the forces to initiate and propagate leg swing at about 10% of the total energy cost of walking [8]. To maintain stability, the center of mass must be controlled relative to the base of support in anterior-posterior and medio-lateral directions [11, 9]. It has been reported that specifically medio-lateral stability is under active control [11, 9], and thus entails an energetic cost [9]. The energy cost used to maintain medio-lateral gait stability can increase in patients with neurological disorders [5, 6]. However, to date, it is unclear how much energy exactly is required for the control of medio-lateral stability.

Previous studies have experimentally investigated the energy costs required for the control of medio-lateral stability [12, 13, 9, 14–17, 10], using a spring-like construction that externally stabilizes walking in the medio-lateral direction (see **Figure 1**). However, results were inconsistent [12, 13, 9, 14–17, 10]. For example, on the one hand, Donelan et al. [9] reported a significant reduction in energy cost when walking with external lateral stabilization at both preferred and zero step widths. While on the other hand, Dean et al. [13] did find a significant reduction in energy cost only when walking at zero step width. There may be several reasons for contradictory results. Differences in task constraints, such as walking at zero or preferred step width [13, 9], walking with or without arm swing [10], and differences in habituation time may have influenced effects of external lateral stabilization on energy cost. In addition, the effect of external lateral stabilization on energy cost may have been influenced by differences in stabilization devices, such as fixed or movable springs in anterior-posterior direction [18], differences in spring stiffness [9, 14, 10] and in rope lengths used to attach the bilateral springs to the participant. For instance, the springs fixed in anterior-posterior direction used in previous studies [12, 13, 9, 10] may provide unwanted assistance in the anterior direction and assist the propulsive force generated by the participants (cf. [18]). Although some studies tried to minimize these potential unwanted effects by using long ropes to connect the springs to the participants (i.e. 8.5 and 14.5 m [9, 10], respectively), other studies with shorter ropes (i.e. 3.0 and 4.0 m [12, 13], respectively) may have overestimated the energy cost savings of external lateral stabilization. Finally, the effect of external lateral stabilization on energy cost may have been missed in some studies due to low statistical power, as studies had relatively small sample sizes (between 8-17 participants) [19, 20].

**Figure 1.**
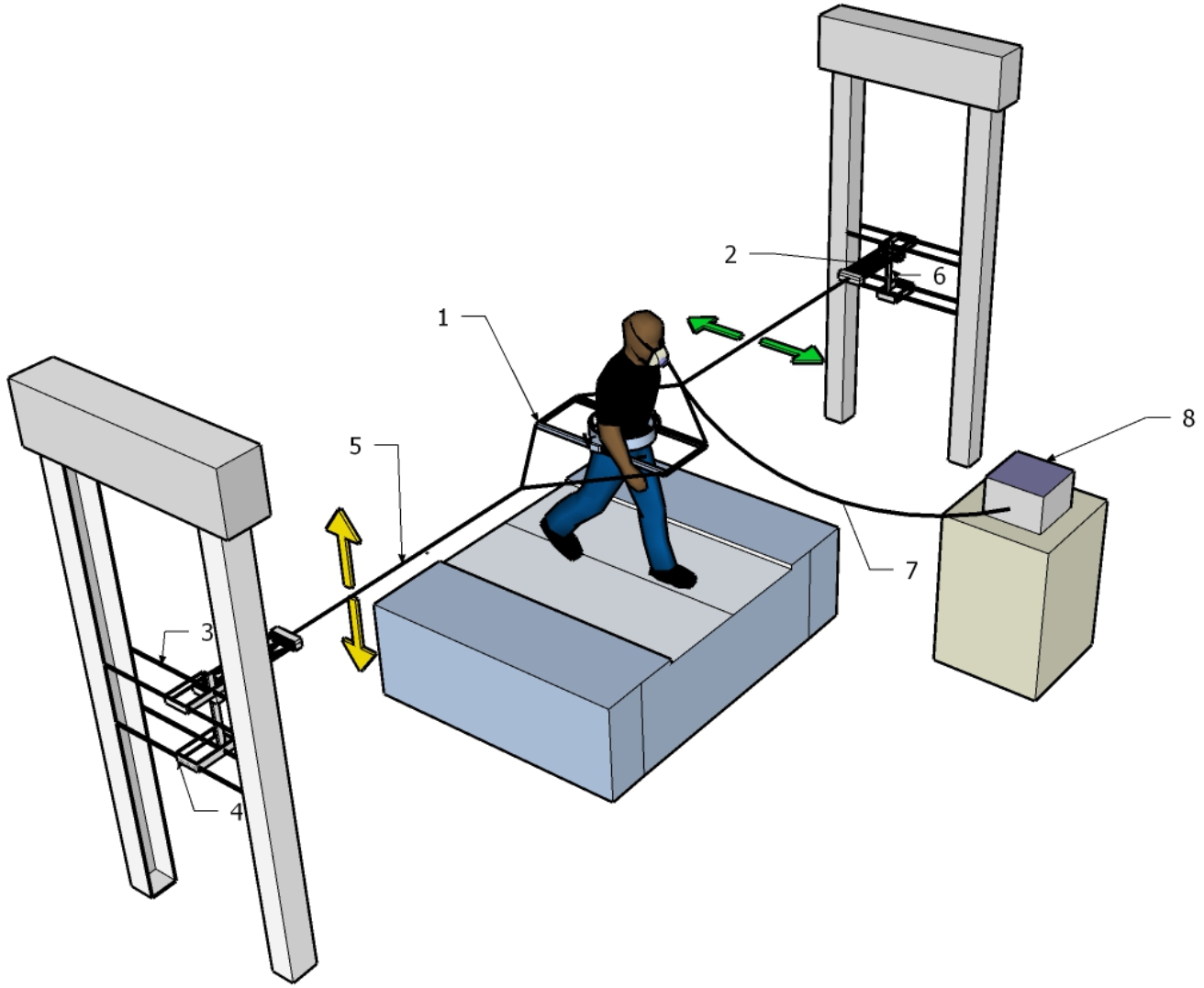
Schematic representation of an external lateral stabilization device used by [16]; (1) Frame; (2) springs; (3) height-adjustable horizontal rail; (4) ball-bearing trolley freely moving in anterior-posterior direction (green arrows show the degree of freedom in anterior-posterior direction); (5) rope attached to frame; (6) vertical rail (yellow arrows show the degree of freedom in vertical direction); (7) sample line and (8) gas analyzer system.

Conducting a meta-analysis on data from the previously published studies that takes the aforementioned differences in task constraints and experimental devices into account may provide a more precise estimate of the true effect of external lateral stabilization on the energy cost of walking. Such analyses may also provide recommendations for future studies using external stabilization. Therefore, the aim of the current systematic literature review with meta-analysis was to provide a synthesis of the findings of previous studies in which the effect of external lateral stabilization on energy cost of walking in healthy young adult participants was investigated. To do so, we explored whether experimental design factors (i.e. step width, arm swing, rope length, spring stiffness, and habituation time) affected the difference in energy cost between walking with and without external lateral stabilization.

## 2. Methods

### 2.1. Search strategy

The Preferred Reporting Items for Systematic Reviews and Meta-Analysis (*PRISMA*)-statement (see: www.prisma-statement.org) was followed as our review protocol. A comprehensive search was performed in PubMed, Google Scholar, and Cochrane library databases. These databases were searched from their commencements up to 19 Feb 2021. The following terms (including synonyms and closely related words) were searched as index terms or keywords without date, language and publication status restrictions: “external lateral stabilization”, “energy cost”, “gait”, “walking”. The full search strategy with the query terms used for each of the three databases is detailed in Supplementary materials. Title, Keywords and Abstract were evaluated by the first author to select the initial list of the papers. A further selection was based on the methodology of the studies and was conducted by the first author.

### 2.2. Inclusion and exclusion criteria

We only included studies published in peer reviewed journals that reported and compared net energy cost of walking with and without external lateral stabilization. We considered those studies which included a group of healthy young adult participants. Young adult participants were defined as having a mean age between 18-41 years as suggested by [4]. We excluded one study [10] from our meta-regression because the individual participant data of this study were not accessible. However, this study was included in our systematic review. All studies which included participants with disabilities, diseases, and injuries only [5, 6] were excluded.

### 2.3. Data extraction strategy

We requested the individual participant data on energy cost from included studies. From the data provided and the data collected in our laboratory, we extracted all the data on net energy cost of walking with different walking speeds, spring stiffnesses and walking patterns (i.e. walking at preferred and zero step widths as well as walking with and without arm swing). If the unit in which net energy cost was reported was not in the SI unit of J kg^-1^ m^-1^, we converted the reported values accordingly.

### 2.4. Overview of analysis

All the data and code used to statistically summarize the effect of external lateral stabilization on net energy cost (i.e. Stabilization effect) can be found at https://surfdrive.surf.nl/files/index.php/s/nawqssParn7dW6D. For our analyses, we considered the potential stabilization device and design factors, which might influence effects of external lateral stabilization on net energy cost.

### 2.5. Effect size calculation

To improve comparability of the effect size of Stabilization between studies, we first performed a repeated measure ANOVA for each study from which we calculated 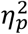, which indicates the sum squares (SS) of the Stabilization effect in relation to the summation of the sum squares of the Stabilization effect and the sum squares of the error associated with the Stabilization effect:

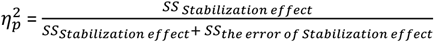

### 2.6. Meta-regression analysis

We performed multilevel analyses on the data to investigate whether Step width, Arm swing, Rope length, Spring stiffness, and Habituation time affect the difference in energy cost between walking with and without external lateral stabilization. Net energy cost_*Stabilized*_ – Net energy cost_*Non-Stabilized*_ was thus calculated as outcome variable and each of the aforementioned design factors were considered as predictor variable. Predictor variables were either continuous or categorical with reference categories as indicated in **Table 1**.

**Table 1:** Predictor variables

| Predictor variable | Category (for categorical predictor variables) or unit of measurement (range) (for continuous predictor variables) |
| --- | --- |
| Step width | Preferred step width (reference) or zero step width |
| Arm swing | With arm swing (reference) or without arm swing |
| Rope length | 1/rope length (m) (range 0-0.33) |
| Spring stiffness | N/m (range 760 -2200 N/m) |
| Habituation time | Minutes (range 0 -7.5 minutes) |

The outcome and predictor variables were clustered in a hierarchical structure. We nested these variables under three Levels: the Participant (Level 1), the Study to which the participant belonged (Level 2) and the Laboratory in which the study was conducted (Level 3) (**Figure 2**). Using the aforementioned levels and predictor variables, we performed multilevel analyses in which random intercepts and random slopes were incorporated into the regression models to attribute the variation in values of the outcome variable to the relevant levels and predictor variables. Analyses started with a model in which each of the design factors separately was considered as a predictor of the outcome variable including a random intercept for Participant (accounting for the repeated measures within participants because of the different conditions (levels of design factors) at which participants had to walk within the experimental studies). To explore whether a random slope in addition to the random intercept for Participant would improve the regression model, the model with random intercept and slope was compared to the model with random intercept only. Compared to the model with random intercept only, the model with random intercept and random slope was preferred if the Bayesian Information Criterion (BIC) decreased and also if the change in −2LL (i.e. −2 x log-likelihood) value between two models was significant (α < 0.05) [21]. The model that resulted from this initial analysis was subsequently hierarchically explored for additional random intercepts and slopes for Levels 2 (Study) and 3 (Laboratory), again using BIC and −2LL values (α < 0.05) [21]. In addition to the simple regression models with only a single predictor, the combined effect of the predictors was also investigated using multiple regression. For combinations of predictor variables that correlated r > 0.70 among each other, the regression analyses were not performed to avoid invalid results due to collinearity. The multilevel analyses were performed with the nlme package [22] in Rstudio [23] software. The R code and the results can be explored in more detail in the supplemental materials (**Tables S3–S10**).

**Figure 2.**
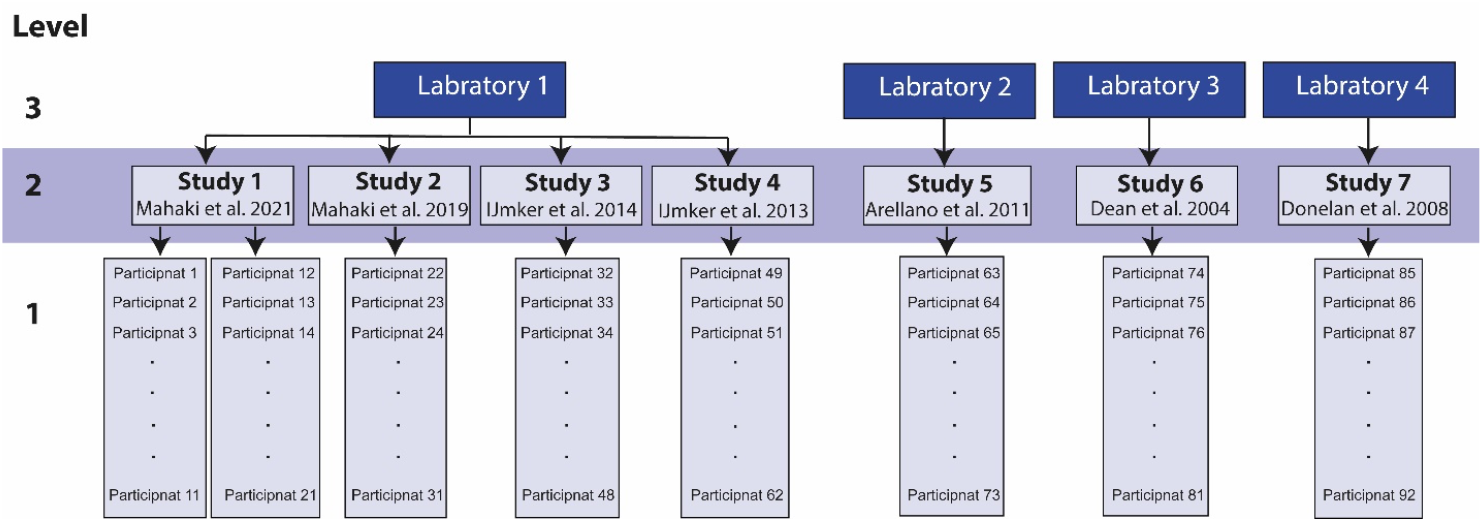
Data layout for multilevel modeling.

## 3. Results

### 3.1. Studies characteristics

We included eight studies (**Figure 3** and **Table 2**), with a range of 8-17 participants per study, reporting on a total of 104 participants. Walking trials of five [14–17] or seven [12, 13, 9, 10] minutes were used in these studies. The last two [14, 15] or three [12, 13, 9, 16, 17, 10] minutes of these walking trials were selected to calculate net energy cost. Stabilized walking was studied at several walking speeds (mean 1.10 m/s, range 0.83-1.70 m/s). Most studies [12, 1, 14–17], but not all [13], randomized the within participant order of non-stabilized and stabilized walking among the participants. In most studies, participants had to walk with restricted transverse and frontal plane pelvis rotations as well as restricted vertical pelvis displacement [12, 13, 9, 14, 15, 10], and had to hold their arms crossed across their chest in stabilized walking (to avoid contacting the external stabilizer) [12, 13, 9]. In some studies, however, normal transverse [14–17] and frontal [17] plane pelvis rotations, vertical pelvis displacement [16] as well as arm swing [14–17, 10] were allowed in stabilized condition. The effects of external lateral stabilization were tested when walking with preferred step width [13, 9, 14–17, 10] and with zero step width [12, 13, 9]. The habituation time (i.e. exposure to the external lateral stabilization before measurements started) varied from 0 [9, 14] to 7.5 [12, 13, 10] min.

**Figure 3.**
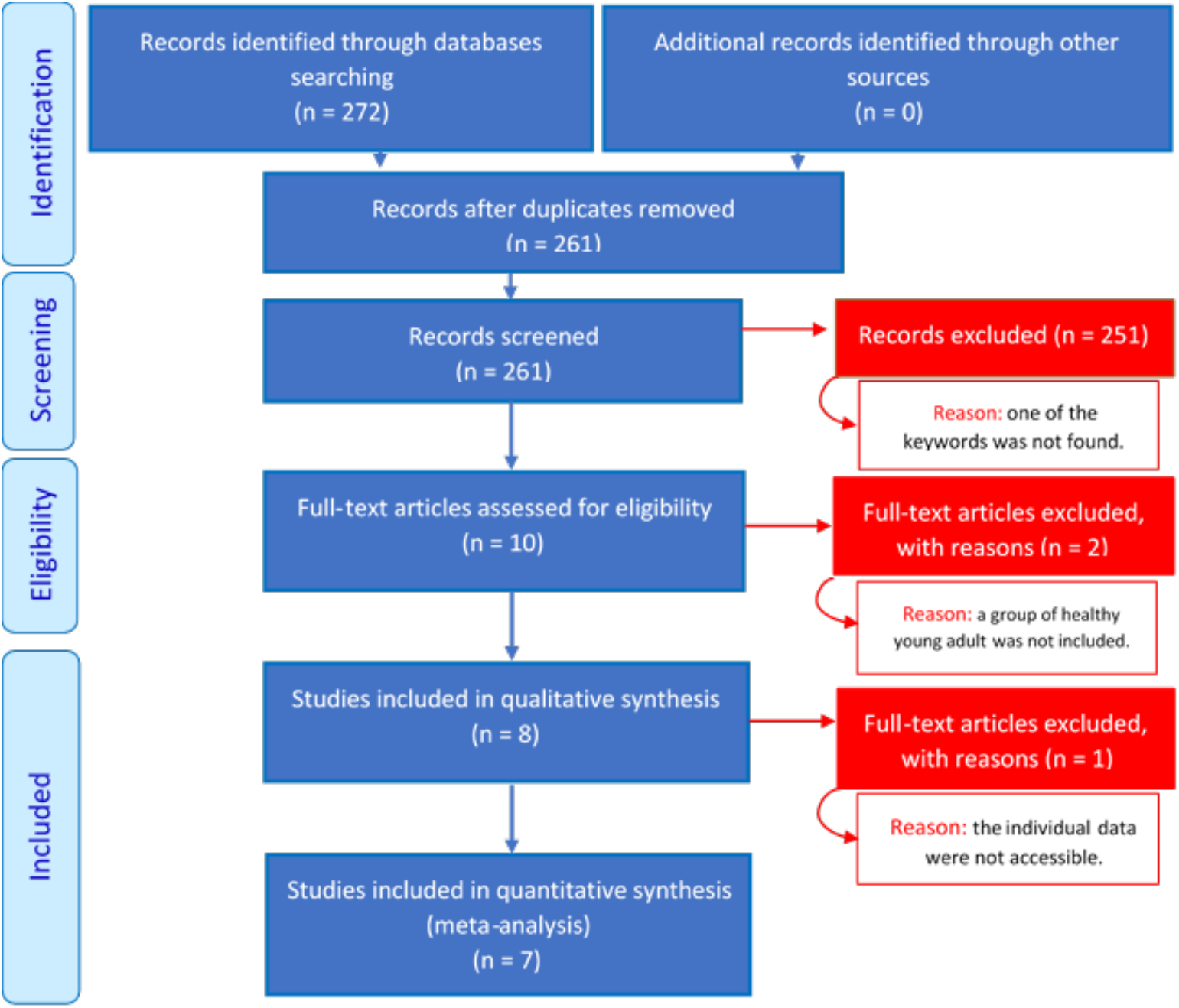
PRISMA flow diagram of the search and selection strategy.

**Table 2.**
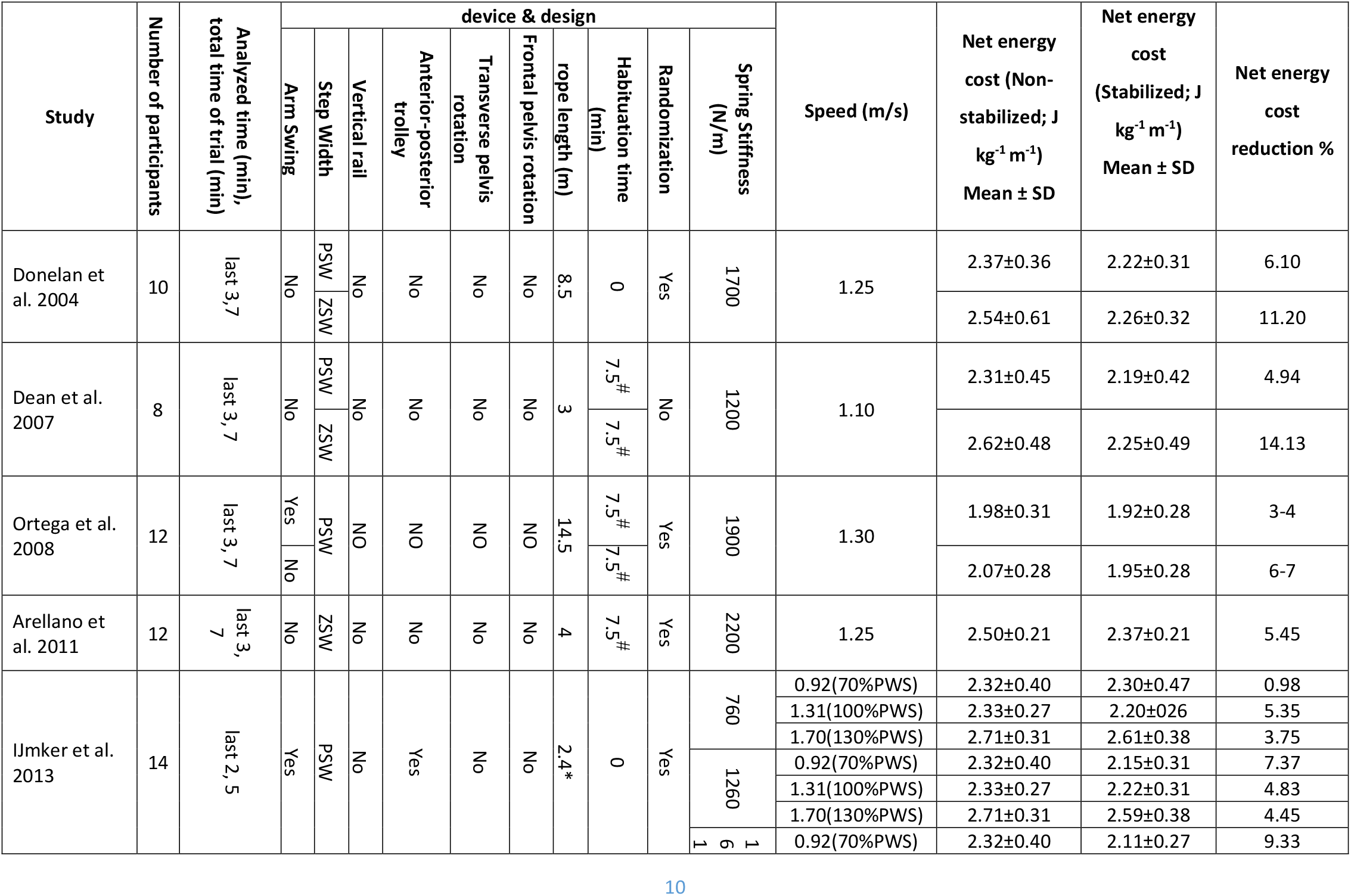
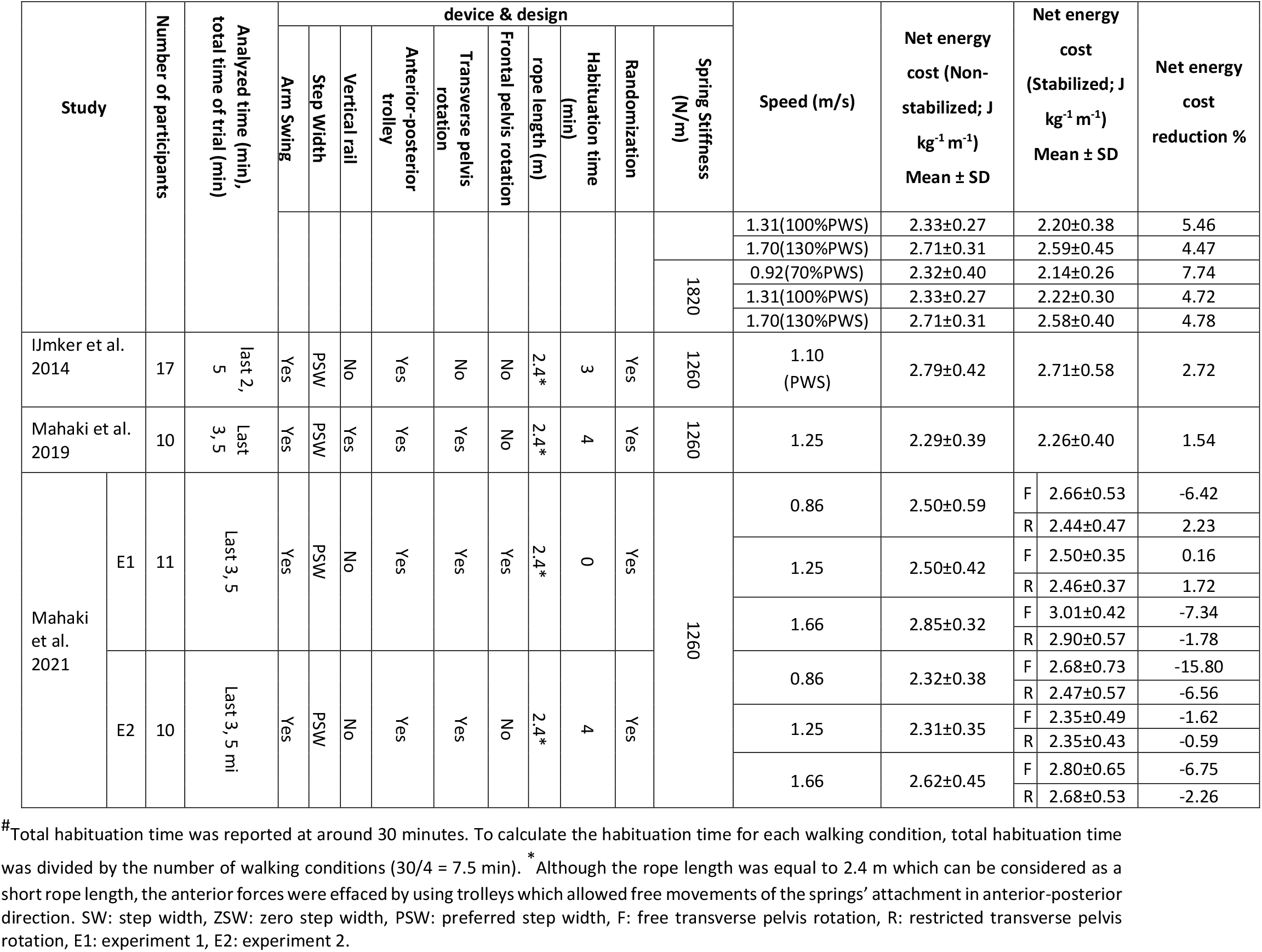
Summary of studies that assessed effects of external lateral stabilization on energy cost in healthy young adult participants.

Regarding the stabilization device, there were considerable differences in the stiffness of the springs used (mean ~1400 N/m, range 760-2200 N/m). Also, the anterior-posterior drift of participants was restricted by the fixed springs in anterior-posterior direction in some studies [12, 13, 9, 10]. To minimize this effect, some studies used long ropes to connect the springs to the participants (i.e. 8.5 and 14.5 m [9, 10], respectively), while other studies used shorter ropes (i.e. 3.0 and 4.0 m [12, 13], respectively). In other studies [14–17], participants could freely move in anterior-posterior direction since the bilateral springs were attached to trolleys.

In non-stabilized walking at preferred and zero step widths, the net energy costs (n = 92) were 2.49±0.42 and 2.55±0.44 J kg^-1^ m^-1^ (mean ± s.d.), respectively. In stabilized walking (n = 92) at preferred and zero step widths the energy costs were 2.44±0.48 and 2.30±0.33 J kg^-1^ m^-1^ (mean ± s.d.), respectively. The relative reductions in net energy cost in stabilized walking at preferred and zero step widths were 2.01% and 9.08%, respectively.

#### Individual study results

The included studies showed mixed findings on the effect and effect size of external lateral stabilization on energy cost of walking (**Figures 4A & B**).

**Figure 4.**
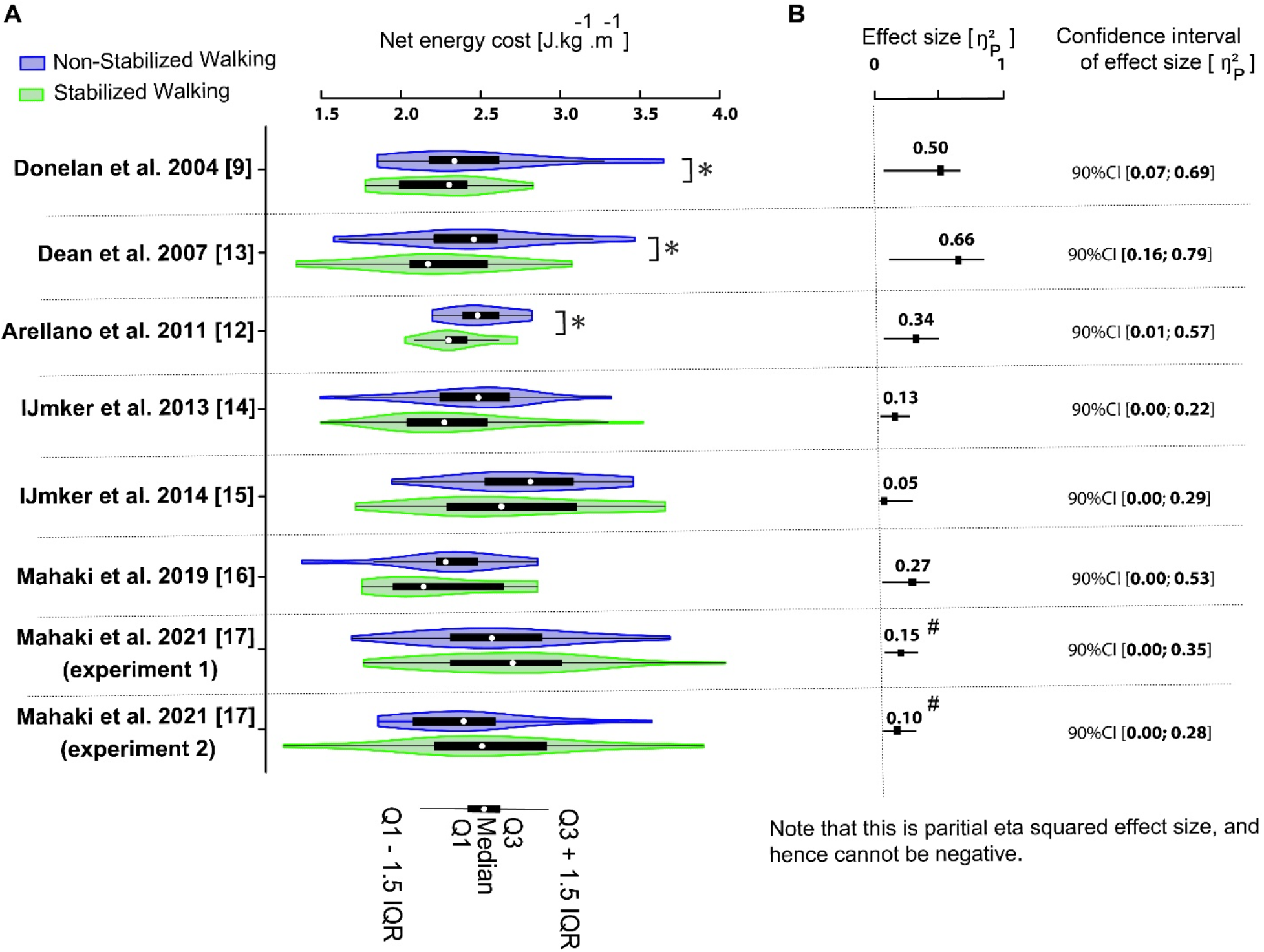
**(A)** The effect of external lateral stabilization on energy cost of walking in healthy young participants. Available data were averaged over any conditions that were non-stabilized and stabilized (e.g step width, arm swing, spring stiffness). **(B)** The size and confidence interval of Stabilization effect on energy cost for each study. * denotes a significant reduction in energy cost by stabilization based on statistical analyses performed on data provided (see Supplementary material, Table S1). # indicates that the effect is opposite (i.e. stabilized walking costs more energy), which cannot be appreciated from the effect size itself, as it’s a squared metric.

Several studies found significant reductions in energy cost due to walking with external lateral stabilization. Donelan et al. [9] reported a significant 5.7% and 9.2% reduction in energy cost during preferred and zero step width conditions, respectively. Arellano et al. [12] reported a significant 5.5% reduction in energy cost at zero step width. In contrast, Dean et al. [13] did not find a significant reduction in energy cost at preferred step width, although they did find an effect at a prescribed zero step width condition. Ortega et al.^1^ [10]. reported that with arm swing the effect of lateral stabilization on energy cost was slightly lower (a significant 3-4% reduction), compared to walking without arm swing (a significant 6-7% reduction). Using bilateral springs with different stiffnesses (760, 1260, 1610, and 1820 N/m) on a slider, IJmker et al. [14] reported no significant reduction of energy cost in stabilized conditions. After visual inspection of the data, IJmker et al. [14] found an outlier (i.e., a participant) in their data. They argued that a potential resistance of this participant against the bilateral spring forces led to increased energy cost in stabilized walking, and therefore favored exclusion of the outlier. After the removal of this outlier, IJmker et al. reported a significant reduction of energy cost when the spring stiffness reached 1260 N/m, without any further reduction of energy cost for higher stiffnesses (i.e. 1610 and 1820 N/m). In a second study by IJmker et al. [15], the effect of stabilization failed to reach significance. In two studies by Mahaki et al. [16, 17] with a slightly different stabilization device than IJmker et al. [14, 15] (i.e. two transverse sliders between waist belt and frame, which allowed transverse pelvis rotation), the effect of stabilization again failed to reach significance.

### 3.2. Meta-regression results

For Step width, Arm swing, and Rope length, there was a significant variation when only intercept was allowed to vary across Participants (Level 1: i). However, including Studies (Level 2) and Laboratories (Level 3) did not significantly improve the models (for intercepts and slopes; see Supplementary materials, **Tables S3–S8**). Table 3 shows the univariate regression models with random intercept across Participants (*b_i_*) and each experimental design factor is included separately as the predictor variable of net energy cost difference between stabilized and non-stabilized walking:

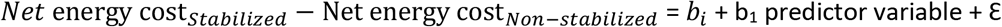

**Table 3.** The univariate analyses of Step width, Arm swing, Rope length, Spring stiffness and Habituation time effects on net energy cost reduction due to walking with external stabilization.

| Predictor variables | Coefficients | value | SE | DF | t-value | p |
| --- | --- | --- | --- | --- | --- | --- |
| Step width | $b_i$ | -0.04 | 0.03 | 266 | -1.08 | 0.28 |
| | $b_1$ | -0.19 | 0.06 | 266 | -3.15 | **0.002** |
| Arm swing | $b_i$ | -0.02 | 0.04 | 267 | -0.60 | 0.55 |
| | $b_1$ | -0.18 | 0.07 | 86 | -2.64 | **0.001** |
| Rope length | $b_i$ | -0.03 | 0.04 | 267 | -0.87 | 0.39 |
| | $b_1$ | -0.64 | 0.27 | 86 | -2.39 | **0.02** |

<sup>1</sup> This study was excluded from our meta-regression analysis because the individual participant data of this study were not accessible.

|  |  |  |  |  |  |  |
| --- | --- | --- | --- | --- | --- | --- |
| Spring stiffness | $b_i$ | 0.03 | 0.07 | 266 | 0.47 | 0.64 |
| | $b_1$ | <-0.001 | <0.001 | 266 | -1.72 | 0.09 |
| Habituation time | $b_i$ | -0.05 | 0.04 | 267 | -1.32 | 0.19 |
| | $b_1$ | -0.01 | 0.01 | 86 | -1.04 | 0.30 |

For walking at zero step width; Step width = 1 and for walking at preferred step width; Step width = 0.
For walking without arm swing; Arm swing = 1 and for walking with arm swing; Arm swing = 0.
$b_i$ represents a significant variation of intercept across Participants (Level 1 = i).

The effect of external lateral stabilization on energy cost of walking was dependent on whether participants walked at preferred or zero step width, as indicated by the significant regression coefficient of Step width (**Table 3**; b_1_). For walking at preferred step width, lateral stabilization did not significantly reduce energy cost (Net energy cost_*Stabilized*_ – Net energy cost_*Non–stabilized*_ = −0.04 J kg^-1^ m^-1^, t(266) = −1.08, p = 0.28) (**Figure 5A**). Rerunning the same regression analyses after changing the reference category of step width from preferred step width to zero step width resulted in a significant intercept of −0.23 J kg^-1^ m^-1^ (i.e., −0.04-0.19, t(266) = −3.15, p = 0.002). These results indicated that the energy cost of stabilized walking at zero step width was 0.23 J kg^-1^ m^-1^ lower than the energy cost of non-stabilized walking at zero step width.

**Figure 5.**
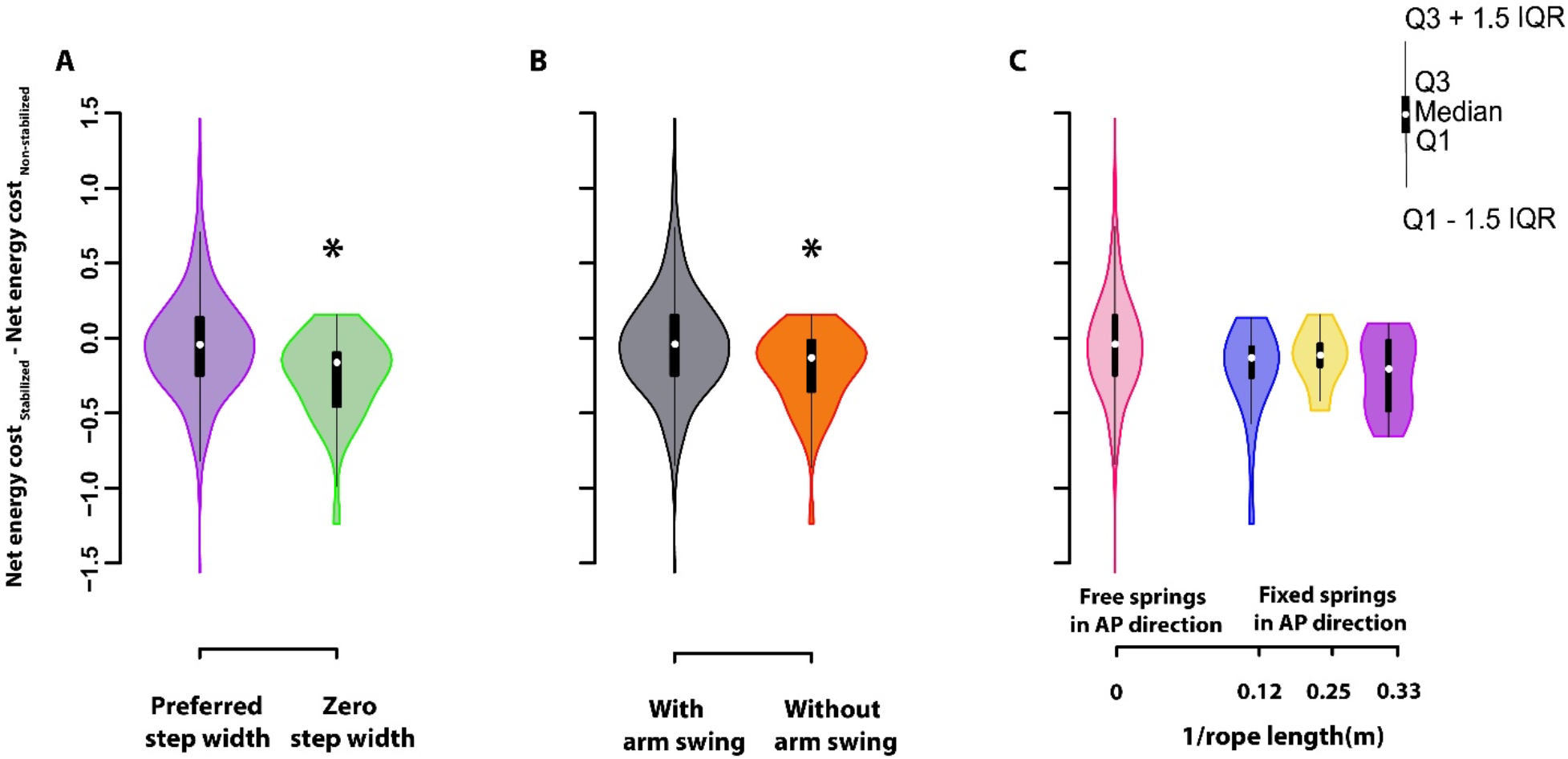
The effect of Step width (**A**), Arm swing (**B**) and Rope length (**C**) on the energy cost reduction due to walking with external lateral stabilization. * denotes a significant reduction in energy cost. Note that the x-axis in 5c represents 1/rope length to include the data for springs that can freely move in anterior-posterior (AP) direction, which approximates an infinite rope length.

The effect of external lateral stabilization on energy cost of walking was dependent on whether participants walked with or without arm swing, as indicated by the significant regression coefficient of Arm swing (**Table 3**; b_1_). For walking with arm swing, lateral stabilization did not significantly reduce energy cost (Net energy cost_*Stabilized*_ – Net energy cost_*Non–Stabiiized*_ = −0.02 J kg^-1^ m^-1^, t(267) = −0.60, p = 0.55) (**Figure 5B**). Rerunning the same regression analyses after changing the reference category of Arm swing from walking with arm swing to without arm swing resulted in a significant intercept of – 0.20 J kg^-1^ m^-1^ (i.e. −0.02-0.18, t(86) = −2.64, p = 0.001). These results indicated that the energy cost of stabilized walking without arm swing was significantly 0.20 J kg^-1^ m^-1^ lower than the energy cost of non-stabilized walking without arm swing.

Rope length had a significant and negative regression effect on energy cost reduction (**Table 3**; b_1_), indicating that lateral stabilization devices with short rope length increased energy cost reduction (**Figure 5C**). However, our post hoc regression analysis (in which the data with an approximated infinite rope length were excluded) did not show the significant effect of rope length on energy cost reduction (see Supplementary materials, **Table S9**). Finally, our analysis did not confirm that Spring stiffness and Habituation time significantly affected the difference in energy cost between stabilized and non-stabilized walking (**Figure 6 and Table 3**; b_1_).

Our multiple regression results showed that the combination of Step width and Arm swing did not provide significant variations compared to the univariate regression models in which Step width and Arm swing were included separately (**Table S10**). The combination of Step width, Arm swing, and Rope length resulted in a high correlation of −0.85 between Arm swing and Rope length. To avoid invalid results due to collinearity, we did not report the results of this multiple regression analysis.

**Figure 6.**
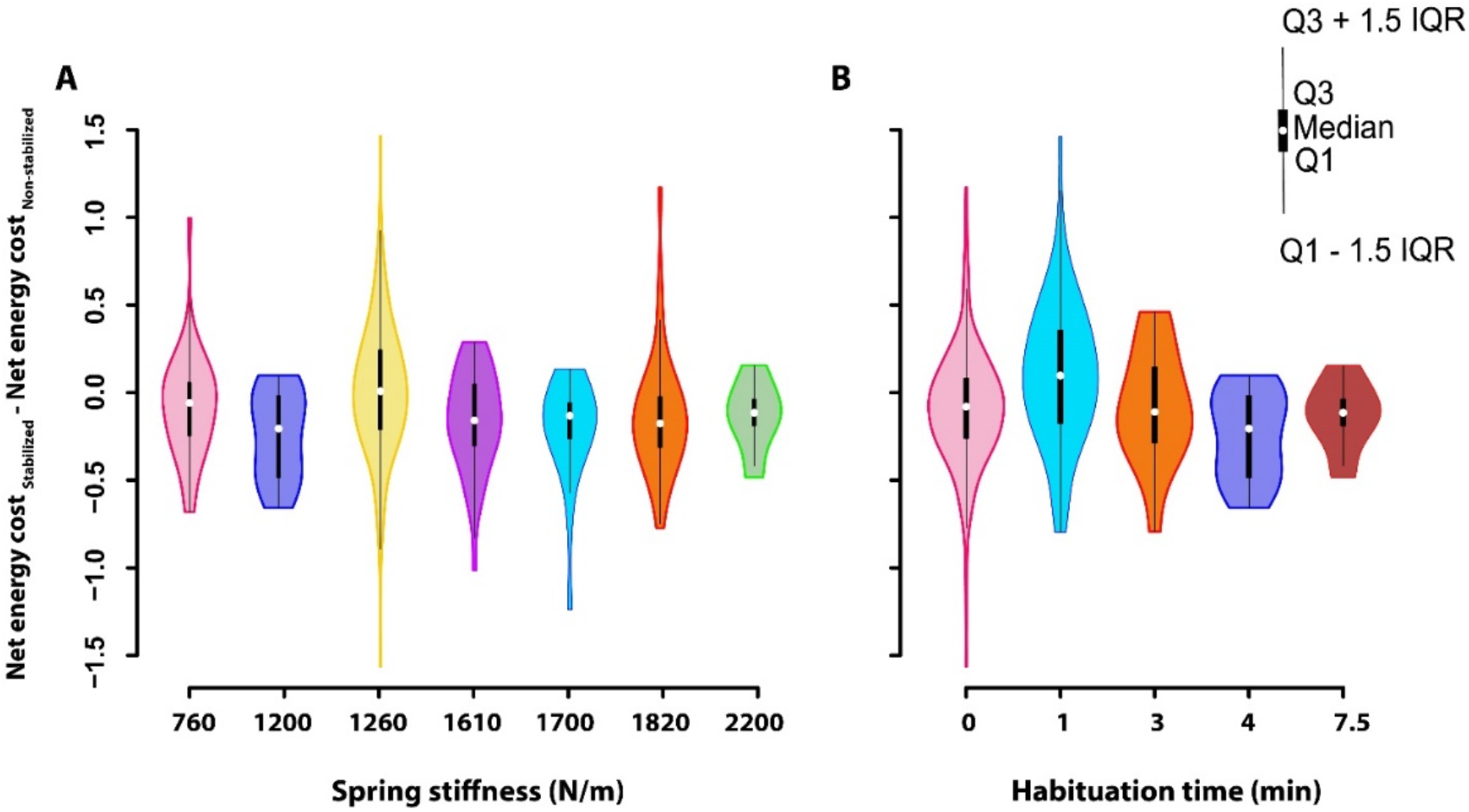
The effect of Spring stiffness (**A**) and Habituation time (**B**) on the energy cost reduction due to walking with external lateral stabilization.

## 4. Discussion

Previous studies reported mixed results on the effects of lateral stabilization on energy cost of walking. To better understand this effect, we performed a systematic review with a meta-analysis of the literature. Our meta-analysis on data from published studies demonstrated no significant variations in energy cost difference between stabilized and non-stabilized walking across studies (Level 2) and laboratories (Level 3). Our results showed that external lateral stabilization reduces energy cost only for walking at zero step width and without arm swing (~8-9%). However, stabilization did not significantly decrease energy cost of walking at preferred step width with or without arm swing (~2-3%). Our meta-analyses also showed that lateral stabilization devices with short rope length increase the energy cost reduction due to walking with external lateral stabilization, while spring stiffness and habituation time do not influence the lateral stabilization effects on energy cost.

There may be several explanations for the different effects of external lateral stabilization on energy cost between walking at preferred and zero step widths. Firstly, the difference in energy cost of (nonstabilized) walking at zero and preferred step width (2.55 vs. 2.49 J kg^-1^ m^-1^) could be due to the use of different stabilizing strategies in these two walking patterns. For walking at preferred step width, foot placement is the main mechanism to control medio-lateral stability [16, 17], while in walking at zero step width, foot placement is constrained and other stabilizing strategies, such as an ankle strategy [24, 25] will be needed. The energy cost of the foot placement is expected to be low, as it needs small modifications in the trajectory of the swing leg only, whereas other strategies involve adaptations of the trajectory of the whole-body mass. Secondly, based on the provided data, in all of the trials related to walking at zero step width, participants also had to walk without arm swing (see **Table 2**). It has been reported that arm swing plays a stabilizing role during walking [26, 10] and this role can become more important when participants have to walk at zero step width. Therefore, due to the constrained foot placement when walking at zero step width and absence of arm swing, maintaining (medio-lateral) stability becomes more challenging [10], which may lead to increased energy cost for control of medio-lateral stability. In line with this, our results showed that for walking at zero step width and for walking without arm swing the effects of lateral stabilization on energy cost reduction was higher than for walking at preferred step width and for walking with arm swing.

It has been reported that the springs fixed in anterior-posterior direction used in previous studies [12, 13, 9, 10] may provide unwanted assistance in the anterior direction and assist the propulsive force generated by the participants (cf. [18]). Humans typically attempt to decrease levels of muscle activation when walking in force fields, also known as “slacking” [27, 28]. Thus, participants may have discovered that walking at the back of the treadmill was less energetically costly with these stabilization devices. Some studies tried to minimize these potential unwanted effects by using long ropes to connect the springs to the participants (i.e. 8.5 and 14.5 m [9, 10], respectively) or by using moveable trolleys to attach springs which prevent such anterior-posterior forces. In line with this, our meta-analysis confirmed that a short rope increases the energy cost reduction due to walking with external lateral stabilization. Although our post hoc analysis did not show the effect of rope length on energy cost reduction in studies with fixed springs in anterior-posterior direction, future lateral stabilization studies are still recommended to use long ropes [9, 10] or trolleys [14–17], to prevent slacking.

Other possible confounders could have been the different spring stiffnesses and habituation times among lateral stabilization studies. However, IJmker et al. [14] reported no significant effect of spring stiffness (760, 1260, 1610, and 1820 N/m). In line with this, our meta-regression analysis confirmed that spring stiffness (range 760-2200 N/m) did not affect the energy cost reduction. Thus, future lateral stabilization studies are recommended to select a spring stiffness between 760-2200 N/m. In addition, habituation time (range 0-7.5 min) did not account for differences in energy cost reduction due to walking with lateral stabilization between studies. One potential reason could be that participants were not fully adapted to stabilized walking after this short period of habituation time. However, all studies reported that they ensured that the metabolic rate had reached a steady state during walking with lateral stabilization, rendering this unlikely.

One of the challenges of a meta-analysis, especially those based on individual data, is that researchers of previously published studies can be unwilling to share their data [29]. However, we were pleasantly surprised by the willingness and openness by which researchers of lateral stabilization studies shared their data with us. By combining the data from our own lab and previously published studies, we performed a meta-analysis, which increased the sample size and thus the power to investigate the lateral stabilization effect on energy cost. Therefore, the current study provides more precise results than each individual study by itself.

We conclude that a small proportion of total energy cost is required to control medio-lateral stability; this proportion is larger when walking with narrow steps and without arm swing. The energy cost reduction due to walking with external lateral stabilization can be influenced by stabilization device properties. For instance, if short ropes are used to connect the bilateral springs to the participant, energy cost reduction can increase due to slacking. However, spring stiffness (range 760-2200 N/m) and habituation time (range 0-7.5 min) do not influence the energy cost reduction with external lateral stabilization.

## Acknowledgements

Sjoerd Bruijn was funded by a VIDI grant (no. 016.Vidi.178.014) from the Dutch Organization for Scientific Research (NWO). We like to thank Trienke IJmker, Jesse Dean, Christopher Arellano and Maxwell Donelan for providing us their data.

## Supplementary materials

### Search Strategy

The full search strategy with the query terms used for each of the three databases are detailed below:

PubMed Session Results (19 Feb 2021) 8 items

((“Energy Metabolism”[Mesh:noexp] OR “Oxidative Phosphorylation”[Mesh] OR energy metabolism[tiab] OR energy expenditure[tiab] OR metabolic cost*[tiab] OR energy cost*[tiab] OR energetic cost*[tiab] OR metabolic consum*[tiab] OR energy consum*[tiab]) AND (“Gait/physiology”[Mesh] OR “Walking/physiology”[Mesh] OR walk[tiab] OR walking[tiab] OR gait[tiab]) AND (“External Lateral Stabilization” [Mesh] OR “Lateral Stabilization” [tiab] OR “Stabilized Walking” [tiab] OR “Stabilized gait”[tiab]))

Google Scholar Session Results (19 Feb 2021) 261 items

(‘energy metabolism’/de OR ‘oxidative phosphorylation’/exp OR ‘energy metabolism’:ab,ti,kw OR ‘energy expenditure’:ab,ti,kw OR

‘metabolic cost*’:ab,ti,kw OR ‘energy cost*’:ab,ti,kw OR ‘energetic cost*’:ab,ti,kw OR ‘metabolic consum*’:ab,ti,kw OR ‘energy

consum*’:ab,ti,kw) AND (‘walking’/exp OR walk:ab,ti,kw OR walking:ab,ti,kw OR gait:ab,ti,kw) AND (‘external lateral stabilization’/de OR lateral stabilization’/exp OR ‘stabilized walking’:ab,ti,kw OR ‘stabilized gait’:ab,ti,kw)

Cochrane Library Session Results (16 Feb 2021) 3 items

((“energy metabolism” or “energy expenditure” or “metabolic cost*” or “energy cost*” or “energetic cost*” or “metabolic consum*” or “energy consum*”) AND (walk or walking or gait) AND (“external lateral stabilization*” or “lateral stabilization*” or “stabilized walking*” or “stabilized gait*”))

### Stabilization effect sizes

We performed repeated measure ANOVA to calculate the Stabilization effect sizes and the confidence interval of the effect size in each study (Table S1). We calculated three types of effect sizes of lateral stabilization studies as reporting multiple effect sizes can yield a greater understanding of a specific effect [1](Table S1). The repeated measure ANOVA together with effect sizes calculation were performed with MATLAB software [2]. We calculated three types of effect sizes: eta squared (*η*^2^), partial eta squared 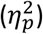 for power analyses, and generalized eta squared 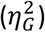 for meta-analyses.

*η*^2^ represents the sum squares of Stabilization effect in relation to the total variance (i.e. SS_total_):

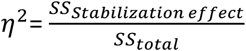

Based on the design of a study, SS_total_ can be calculated differently. For example, in Donelan et al., SS_total_ was calculated as follows:

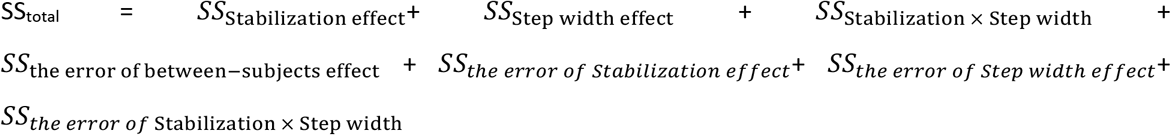

However, in IJmker et al. 2013, SS_total_ was calculated as follows:

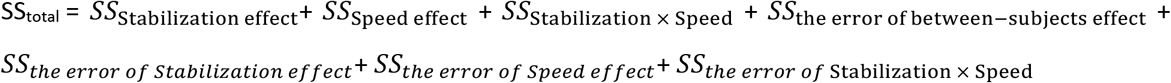

Consequently, *η*^2^ cannot easily be compared between lateral stabilization studies. To improve comparability of effect sizes between studies, we calculated 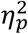 and 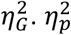 indicates the sum squares of the Stabilization effect in relation to the sum squares of the Stabilization effect and the sum squares of the error associated with the Stabilization effect:

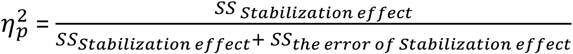

However, Olejnik and Algina [3] reported that the differences in inclusion of covariates or blocking factors between experimental designs (for instance including the age of participations (i.e. young vs old) in the analysis as a between-subjects factor, which will account for some of the variance) can affect the size of 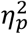. Thus, Olejnik and Algina [3] suggested 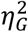 which excludes variation of other factors from effect size calculation and includes variation of individual differences as follows:

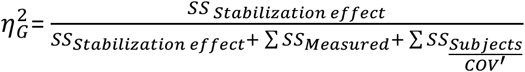

*SS_Measured_* refers to the sums of squares for all the blocking factors or interactions with blocking factors; 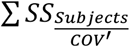 includes the sums of squares involving subjects or covariates (equal to *SS*_total_ in a between-group design). 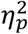 and 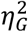 are equal when all factors are manipulated.

**Table S1.**
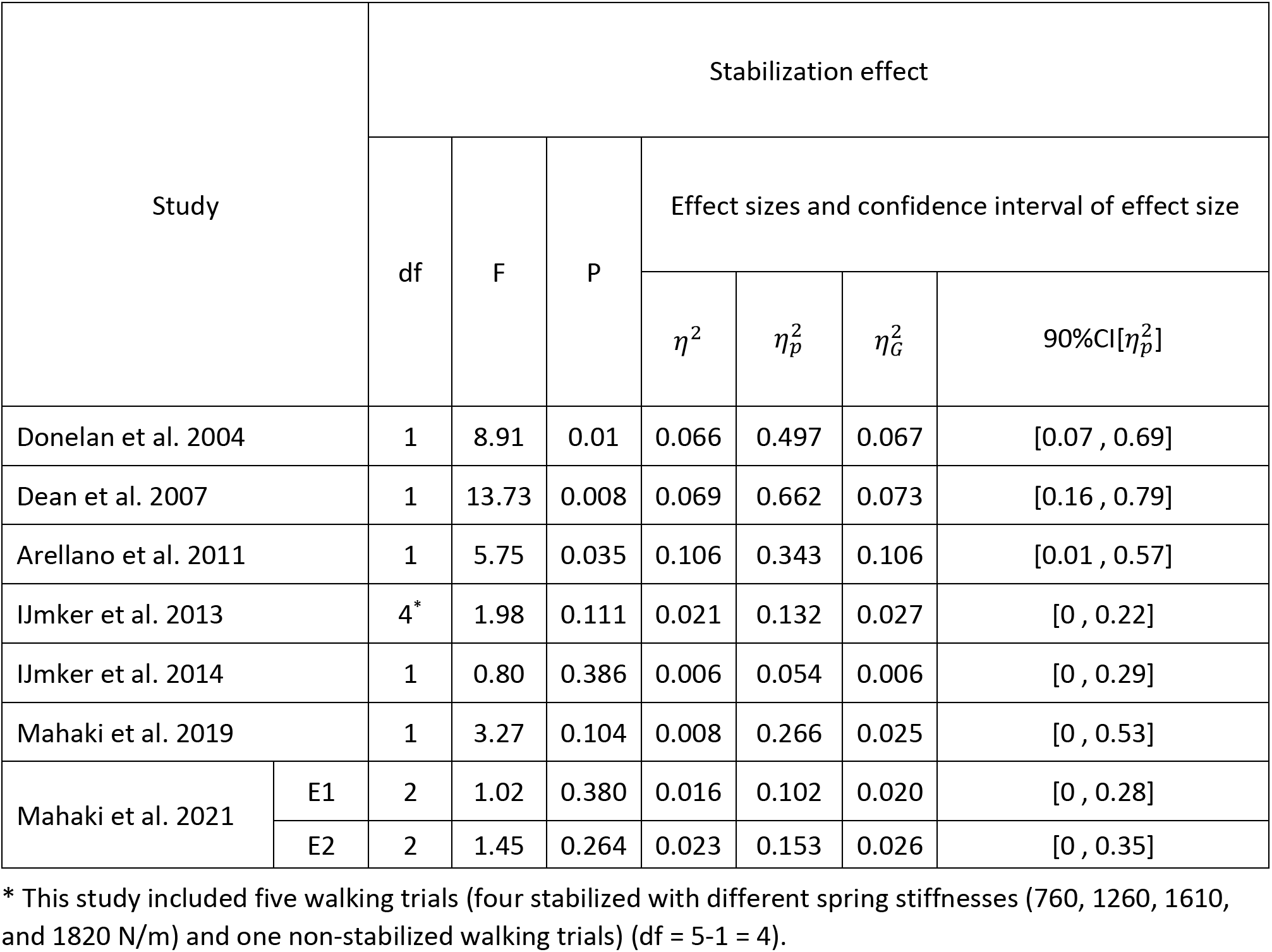
The effect of external lateral stabilization on energy cost of walking

| Study |  | Stabilization effect |  |  |  |  |  |  |
| --- | --- | --- | --- | --- | --- | --- | --- | --- |
|  |  | df | F | P | Effect sizes and confidence interval of effect size |  |  |  |
| | | | | | $\eta^2$ | $\eta_p^2$ | $\eta_G^2$ | 90%CI[$\eta_p^2$] |
| Donelan et al. 2004 |  | 1 | 8.91 | 0.01 | 0.066 | 0.497 | 0.067 | [0.07 , 0.69] |
| Dean et al. 2007 |  | 1 | 13.73 | 0.008 | 0.069 | 0.662 | 0.073 | [0.16 , 0.79] |
| Arellano et al. 2011 |  | 1 | 5.75 | 0.035 | 0.106 | 0.343 | 0.106 | [0.01 , 0.57] |
| Ijmker et al. 2013 |  | 4* | 1.98 | 0.111 | 0.021 | 0.132 | 0.027 | [0 , 0.22] |
| Ijmker et al. 2014 |  | 1 | 0.80 | 0.386 | 0.006 | 0.054 | 0.006 | [0 , 0.29] |
| Mahaki et al. 2019 |  | 1 | 3.27 | 0.104 | 0.008 | 0.266 | 0.025 | [0 , 0.53] |
| Mahaki et al. 2021 | E1 | 2 | 1.02 | 0.380 | 0.016 | 0.102 | 0.020 | [0 , 0.28] |
|  | E2 | 2 | 1.45 | 0.264 | 0.023 | 0.153 | 0.026 | [0 , 0.35] |

\* This study included five walking trials (four stabilized with different spring stiffnesses (760, 1260, 1610, and 1820 N/m) and one non-stabilized walking trials) (df = 5-1 = 4).

### Multilevel analyses

The following model is the model when each design factor (step width, arm swing, rope length, spring stiffness, or habituation time) is included as the predictor of net energy cost difference between stabilized and non-stabilized walking:

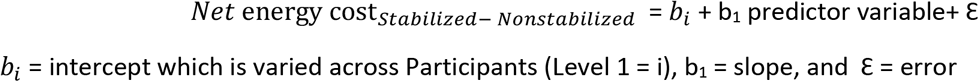

Each of the design factors separately was considered as a predictor of the outcome variable including a random intercept for Participant. To explore whether a random slope in addition to the random intercept for Participant would improve the regression model, the model with random intercept and slope was compared to the model with random intercept only, using ANOVA test. The model with random intercept and random slope was preferred if the Bayesian information criterion (BIC) decreased and also if the change in −2LL (i.e. −2 x log-likelihood) between two models was significant (α < 0.05). The model with random intercept and slope was compared to the model with random intercept only for Level 1(**Table S3**) as follows:

**Table S3.**
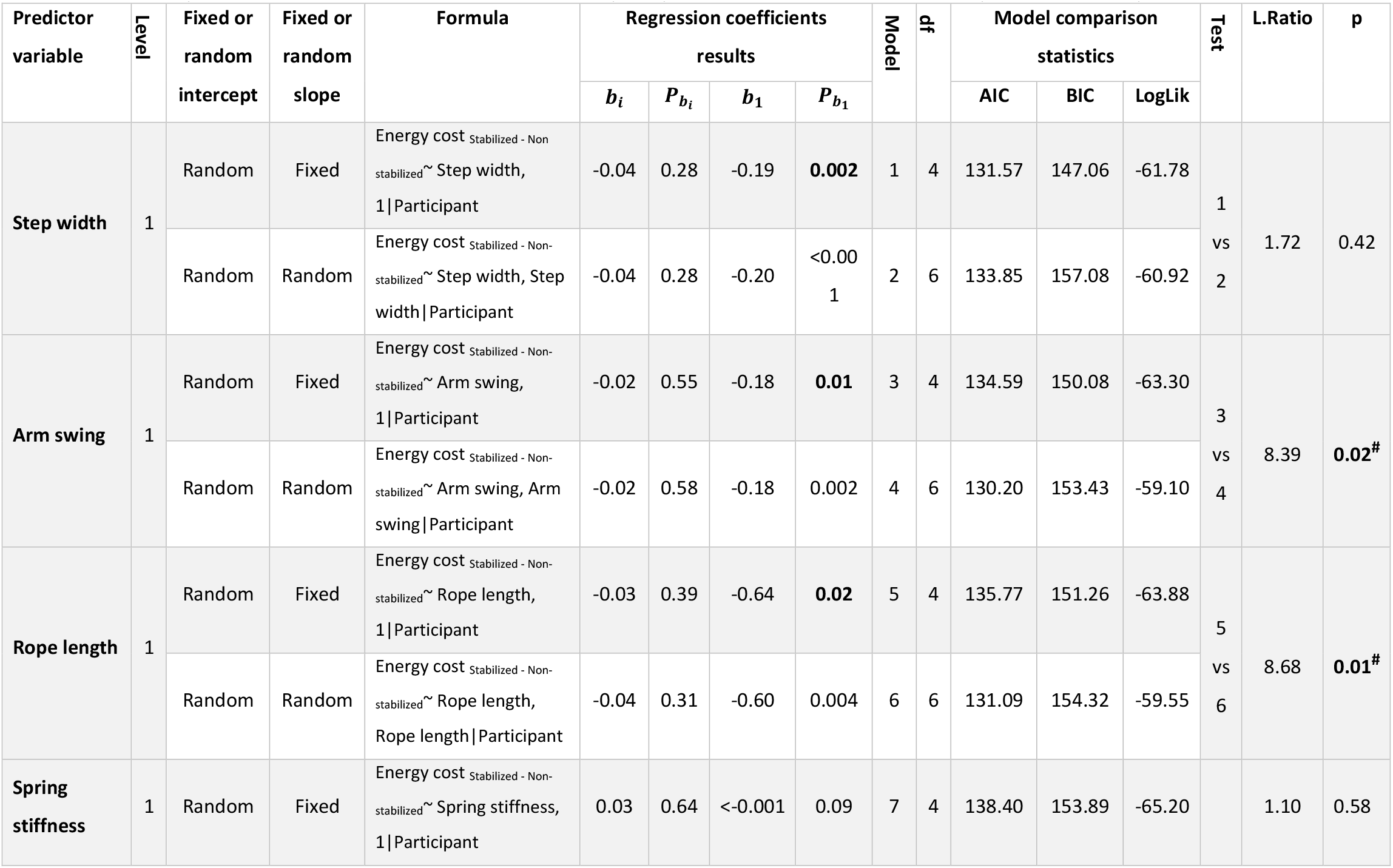
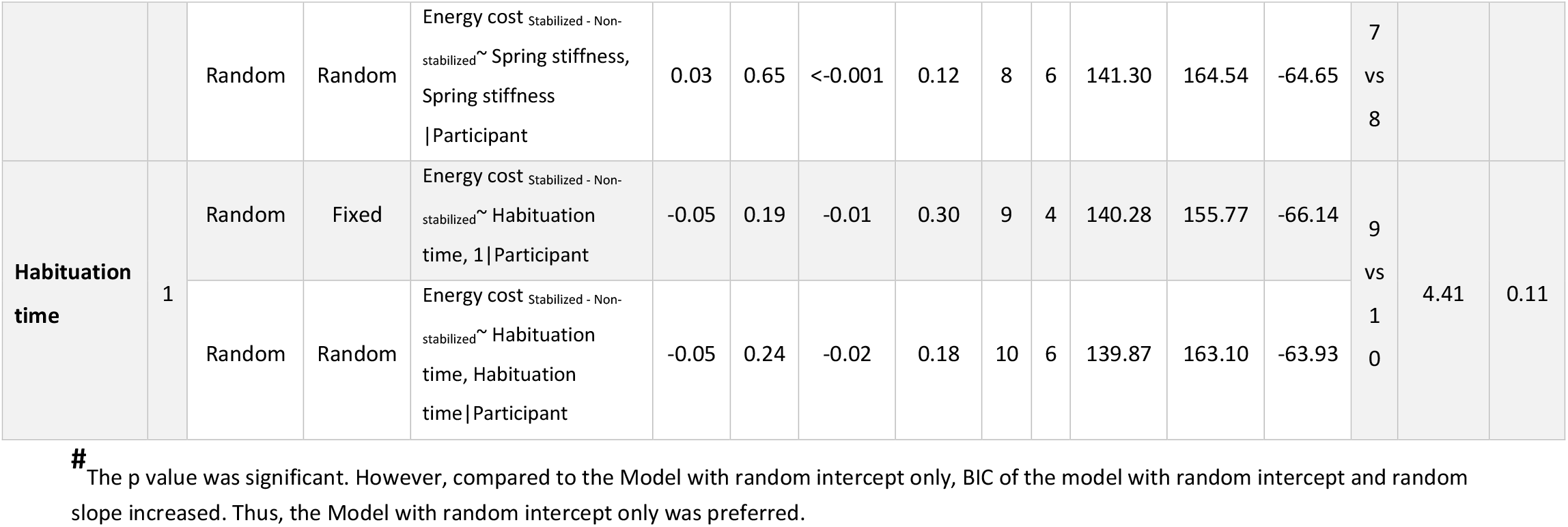
Comparisons between the model with random intercept only and the model with random intercept and random slope (**Level 1**)

For Step width, Arm swing, Rope length, Spring stiffness, and Habituation time, there were significant variations when intercept only was allowed to vary across Participants. For each predictor, the model that resulted from this initial analysis was subsequently hierarchically explored for additional random intercepts and slopes for Levels 2 (Study) as follows (**Table S4**):

**Table S4.** Comparisons between the model with random intercept only and the model with random intercept and random slope (**Level 2**)

| Predictor variable | Level | Fixed or random intercept | Fixed or random slope | Formula | Regression coefficients results |  |  |  | Model | df | Model comparison statistics |  |  | Test | L.Ratio | p |
| --- | --- | --- | --- | --- | --- | --- | --- | --- | --- | --- | --- | --- | --- | --- | --- | --- |
| | | | | | $b_i$ | $P_{b_i}$ | $b_1$ | $P_{b_1}$ | | | AIC | BIC | LogLik | | | |
| Step width | 2 | Random | Fixed | Energy cost Stabilized - Non-stabilized ~ Step width, 1\|Study/Participant | -0.03 | 0.40 | -0.20 | **0.003** | 1 | 5 | 132.98 | 152.34 | -61.49 | 1 vs 2 | 1.78 | 0.78 |
|  |  | Random | Random | Energy cost Stabilized - Non-stabilized ~ Step width, Step width\|Study/Participant | -0.03 | 0.40 | -0.20 | **0.002** | 2 | 9 | 139.21 | 174.06 | -60.60 |  |  |  |
| Arm swing | 2 | Random | Fixed | Energy cost Stabilized - Non-stabilized ~ Arm swing, 1\|Study/Participant | -0.02 | 0.66 | -0.18 | **0.04** | 3 | 5 | 136.34 | 155.71 | -63.17 | 3 vs 4 | 8.15 | 0.09 |
|  |  | Random | Random | Energy cost Stabilized - Non-stabilized ~ Arm swing, Arm swing\|Study/Participant | -0.02 | 0.58 | -0.18 | **0.02** | 4 | 9 | 136.20 | 171.05 | -59.10 |  |  |  |
| Rope length | 2 | Random | Fixed | Energy cost Stabilized - Non-stabilized ~ Rope | -0.03 | 0.51 | -0.65 | 0.07 | 5 | 5 | 137.36 | 156.72 | -63.68 |  | 8.27 | 0.08 |
| | | | | | $b_i$ | $P_{b_i}$ | $b_1$ | $P_{b_1}$ | | | AIC | BIC | LogLik | | | |
|  |  |  |  | length, 1\|Study/<br>Participant |  |  |  |  |  |  |  |  |  | 5<br>vs<br>6 |  |  |
|  |  | Random | Random | Energy cost Stabilized -<br>Non-stabilized~ Rope<br>length, Rope<br>length\|Study/<br>Participant | -0.04 | 0.35 | -0.61 | 0.03 | 6 | 9 | 137.09 | 171.94 | -59.54 |  |  |  |
| Spring stiffness | 2 | Random | Fixed | Energy cost Stabilized -<br>Non-stabilized~ Spring<br>stiffness, 1\|Study/<br>Participant | 0.02 | 0.76 | <-<br>0.00<br>1 | 0.12 | 7 | 5 | 139.14 | 158.50 | -64.57 | 7<br>vs<br>8 | 1.16 | 0.88 |
|  |  | Random | Random | Energy cost Stabilized -<br>Non-stabilized~ Spring<br>stiffness, Spring<br>stiffness \|Study/<br>Participant | 0.03 | 0.76 | <-<br>0.00<br>1 | 0.17 | 8 | 9 | 145.97 | 180.82 | -63.99 |  |  |  |
| Habituation time | 2 | Random | Fixed | Energy cost Stabilized -<br>Non-stabilized~<br>Habituation time, 1\|<br>Study/ Participant | -0.04 | 0.36 | -0.01 | 0.41 | 9 | 5 | 140.71 | 160.07 | -65.36 | 9<br>vs<br>10 | 4.08 | 0.40 |
| | | | | | $b_i$ | $P_{b_i}$ | $b_1$ | $P_{b_1}$ | | | AIC | BIC | LogLik | | | |
|  |  | Random | Random | Energy cost Stabilized - Non-stabilized ~ Habituation time, Habituation time \| Study/ Participant | -0.04 | 0.36 | -0.02 | 0.33 | 10 | 9 | 144.63 | 179.48 | -63.32 |  |  |  |

**Table S6.**
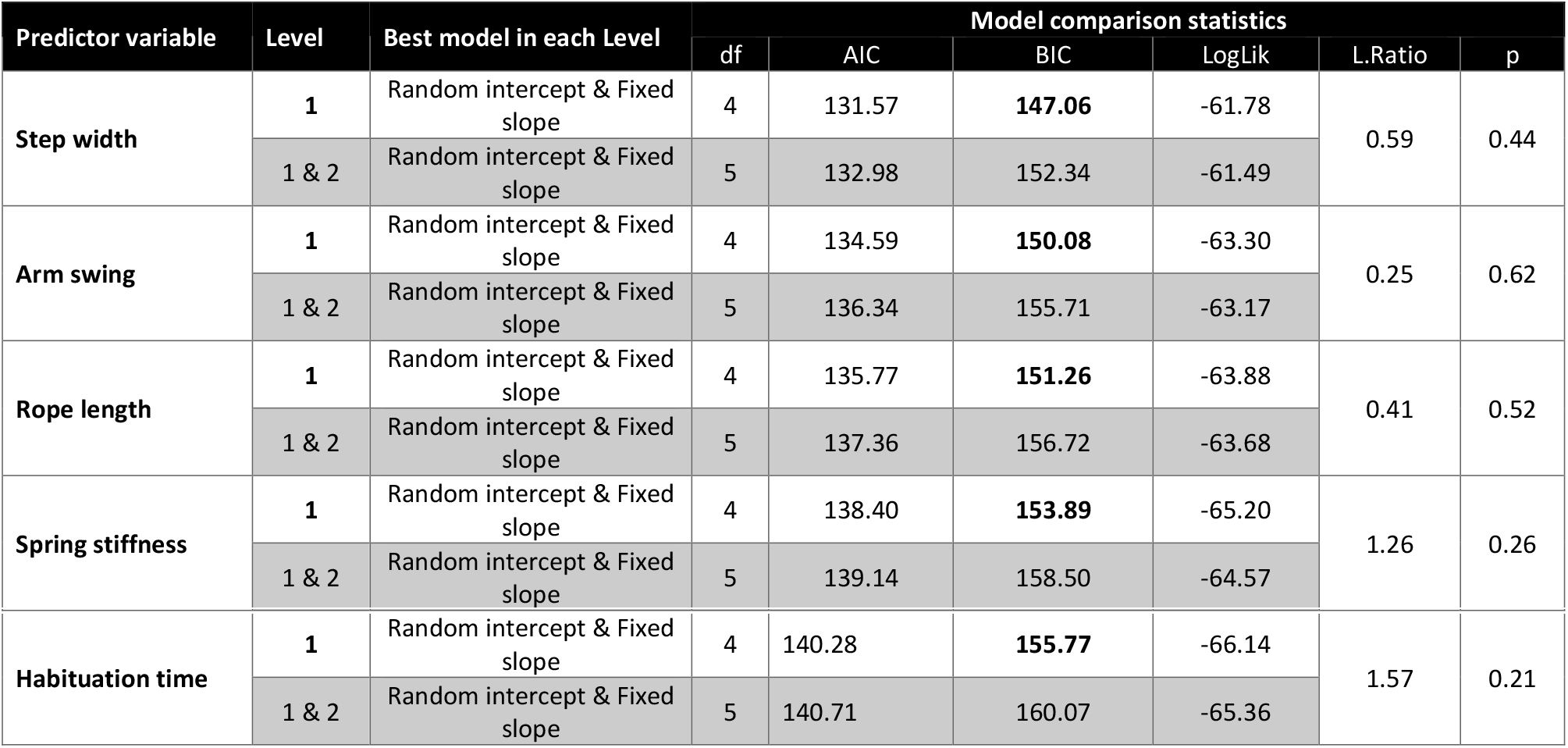
Levels comparison between the best model in <u>Level 1</u> and the best model in <u>Level (1 & 2)</u>.

For Step width, Arm swing, Rope length, Spring stiffness, and Habituation time, there were no significant variations when intercept and slope were allowed to vary across Studies (**Tables S5 and S6**). Thereafter, for each predictor, the model that resulted from the initial analysis (**Table S3**) was subsequently hierarchically explored for additional random intercepts and slopes for Levels 3 (Laboratory) as follows (**Table S7**):

**Table S7.** Comparisons between the model with random intercept only and the model with random intercept and random slope (**Level 3**)

| Predictor variable | Level | Fixed or random intercept | Fixed or random slope | Formula | Regression coefficients results |  |  |  | Model | df | Model comparison statistics |  |  | Test | L.Ratio | p |
| --- | --- | --- | --- | --- | --- | --- | --- | --- | --- | --- | --- | --- | --- | --- | --- | --- |
| | | | | | $b_i$ | $P_{b_i}$ | $b_1$ | $P_{b_1}$ | | | AIC | BIC | LogLik | | | |
| Step width | 3 | Random | Fixed | Energy cost Stabilized - Non-stabilized ~ Step width, 1 \| Laboratory/Participant | -0.04 | 0.28 | -0.19 | 0.002 | 1 | 5 | 133.57 | 152.93 | -61.78 | 1 vs 2 | 1.72 | 0.79 |
|  |  | Random | Random | Energy cost Stabilized - Non-stabilized ~ Step width, Step width \| Laboratory/Participant | -0.04 | 0.28 | -0.20 | <0.001 | 2 | 9 | 139.85 | 174.69 | -60.92 |  |  |  |
| | | | | | $b_i$ | $P_{b_i}$ | $b_1$ | $P_{b_1}$ | | | AIC | BIC | LogLik | | | |
| Arm swing | 3 | Random | Fixed | Energy cost Stabilized - Non-stabilized ~ Arm swing, 1 \| Laboratory/Participant | -0.02 | 0.55 | -0.18 | 0.12 | 3 | 5 | 136.59 | 155.95 | -63.30 | 3 vs 4 | 8.39 | 0.08 |
|  |  | Random | Random | Energy cost Stabilized - Non-stabilized ~ Arm swing, Arm swing \| Laboratory/Participant | -0.02 | 0.58 | -0.18 | 0.08 | 4 | 9 | 136.20 | 171.05 | -59.10 |  |  |  |
| Rope length | 3 | Random | Fixed | Energy cost Stabilized - Non-stabilized ~ Rope length, 1 \| Laboratory/ Participant | -0.03 | 0.39 | -0.64 | 0.14 | 5 | 5 | 137.77 | 157.13 | -63.88 | 5 vs 6 | 8.65 | 0.07 |
|  |  | Random | Random | Energy cost Stabilized - Non-stabilized ~ Rope length, Rope length \| Laboratory / Participant | -0.04 | 0.32 | -0.60 | 0.10 | 6 | 9 | 137.12 | 171.97 | -59.56 |  |  |  |
| Spring stiffness | 3 | Random | Fixed | Energy cost Stabilized - Non-stabilized ~ Spring stiffness, 1 \| Laboratory / Participant | -0.01 | 0.88 | <- 0.001 | 0.14 | 7 | 5 | 139.86 | 159.22 | -64.93 | 7 vs 8 | 1.11 | 0.89 |
|  |  | Random | Random | Energy cost Stabilized - Non-stabilized ~ Spring stiffness, | -0.02 | 0.85 | <- 0.001 | 0.21 | 8 | 9 | 146.75 | 181.60 | -64.38 |  |  |  |
| | | | | | $b_i$ | $P_{b_i}$ | $b_1$ | $P_{b_1}$ | | | AIC | BIC | LogLik | | | |
|  |  |  |  | Spring stiffness \| Laboratory/ Participant |  |  |  |  |  |  |  |  |  |  |  |  |
| Habituation time | 3 | Random | Fixed | Energy cost Stabilized - Non-stabilized ~ Habituation time, 1\| Laboratory / Participant | -0.09 | 0.12 | -0.01 | 0.60 | 9 | 5 | 141.77 | 161.13 | -65.83 | 9 vs 10 | 4.64 | 0.33 |
|  |  | Random | Random | Energy cost Stabilized - Non-stabilized ~ Habituation time, Habituation time\| Laboratory/ Participant | -0.09 | 0.15 | -0.01 | 0.50 | 10 | 9 | 145.13 | 179.98 | -63.56 |  |  |  |

**Table S8.** Levels comparison between the best model in <u>Level 1</u> and the best model in <u>Level (1 & 3)</u>.

| Predictor variable | Level | Best model in each Level | Model comparison statistics |  |  |  |  | p |
| --- | --- | --- | --- | --- | --- | --- | --- | --- |
|  |  |  | df | AIC | BIC | LogLik | L.Ratio |  |
| Step width | 1 | Random intercept & Fixed slope | 4 | 131.57 | **147.06** | -61.78 | 1.26 | 0.99 |
|  | 1 & 3 | Random intercept & Fixed slope | 5 | 133.57 | 152.93 | -61.78 |  |  |
| Arm swing | 1 | Random intercept & Fixed slope | 4 | 134.59 | **150.08** | -63.30 | 1.47 * 10<sup>-7</sup> | 0.99 |
|  | 1 & 3 | Random intercept & Fixed slope | 5 | 136.59 | 155.95 | -63.30 |  |  |
| Rope length | 1 | Random intercept & Fixed slope | 4 | 135.77 | **151.26** | -63.88 | 8.65 | 0.12 |
|  | 1 & 3 | Random intercept & Fixed slope | 5 | 137.12 | 171.97 | -59.56 |  |  |
| Spring stiffness | 1 | Random intercept & Fixed slope | 4 | 138.40 | **153.89** | -65.20 | 0.54 | 0.46 |
|  | 1 & 3 | Random intercept & Fixed slope | 5 | 139.86 | 159.22 | -64.93 |  |  |
| Habituation time | 1 | Random intercept & Fixed slope | 4 | 140.28 | **155.77** | -66.14 | 0.51 | 0.47 |
|  | 1 & 3 | Random intercept & Fixed slope | 5 | 141.77 | 161.13 | -65.83 |  |  |

For Step width, Arm swing, Rope length, Spring stiffness, and Habituation time, there were no significant variations when intercept and slope were allowed to vary across Laboratory (**Tables S7** and **S8**).

**Table S9.**
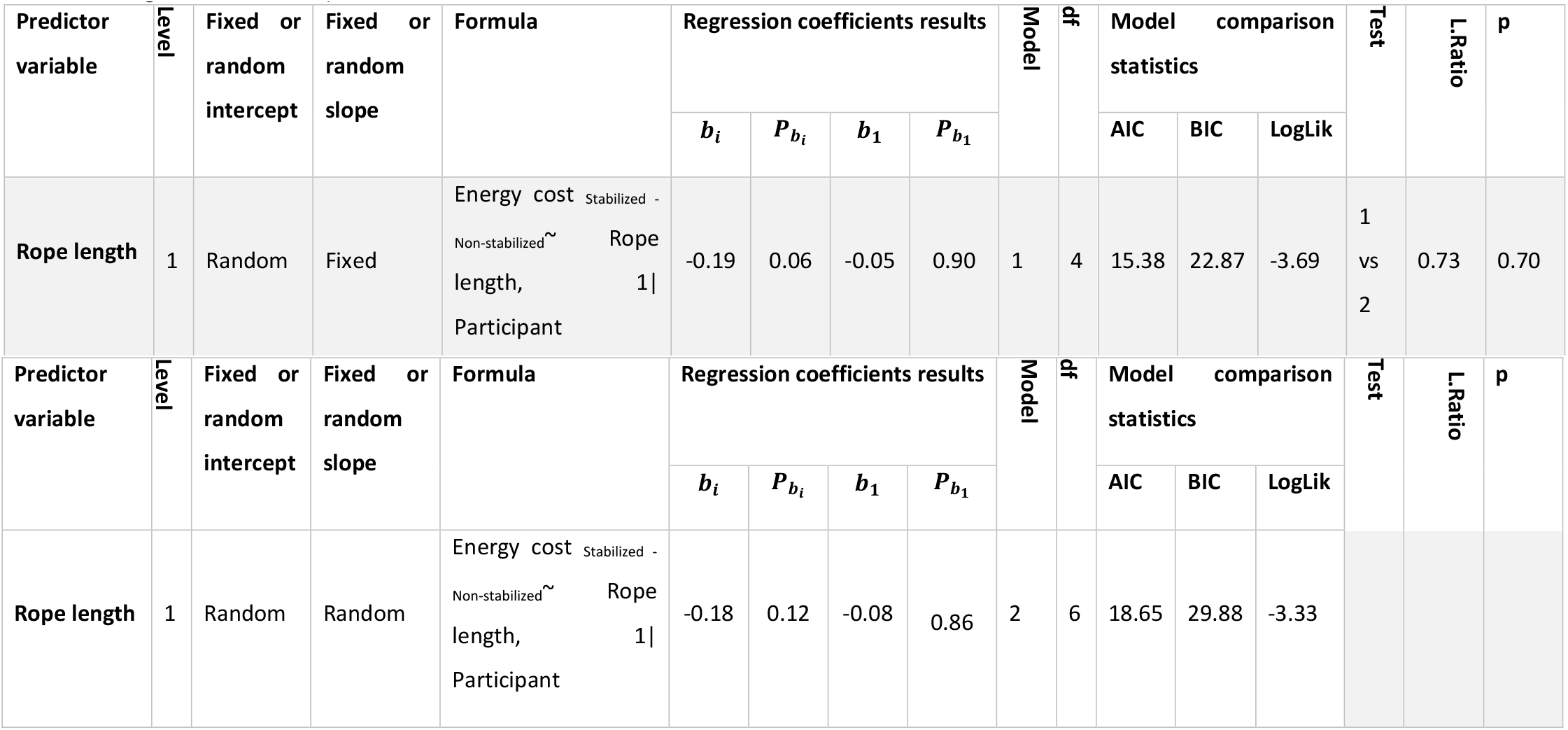
The effect of rope length on energy cost reduction due to walking with lateral stabilization (data with an approximated infinite rope length were excluded)

In addition to the simple regression models with only the design factors as single predictors of outcome variable, the combined effect of the design factors was also investigated using multiple regression (**Table S9**).

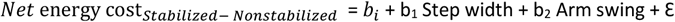

*b_i_* = intercept which is varied across Participants (Level 1 = i), b_1_ = regression coefficient for Step width, b_2_ = regression coefficient for Arm swing and ε = error

For combinations of predictor variables that correlated r > 0.70 among each other, the regression analyses were not performed to avoid invalid results due to collinearity.

**Table S10.** The combined effect of the design factors

| Predictor variable | Level | Fixed or random intercept | Fixed or random slope | Formula | Regression coefficients results |  |  |  |  |  | Model | df | Model comparison statistics |  |  | Test | L.Ratio | p |
| --- | --- | --- | --- | --- | --- | --- | --- | --- | --- | --- | --- | --- | --- | --- | --- | --- | --- | --- |
| | | | | | $b_i$ | $P_{b_i}$ | $b_1$ | $P_{b_1}$ | $b_2$ | $P_{b_2}$ | | | AIC | BIC | LogLik | | | |
| Step width + Arm swing | 1 | Random | For both predictors Fixed | Energy cost Stabilized - Non-stabilized ~ Step width + Arm swing, 1 \| Participant | 0.02 | 0.55 | -0.15 | 0.05 | -0.07 | 0.38 | 1 | 5 | 132.79 | 152.15 | -61.39 | - | - | - |
| Step width | 1 | Random | Fixed | Energy cost Stabilized - Non-stabilized ~ Step width, 1 \| Participant | -0.04 | 0.28 | -0.19 | **0.002** | - | - | 2 | 4 | 131.57 | 147.06 | -61.78 | 1 vs 2 | 0.78 | 0.38 |
| Arm swing | 1 | Random | Fixed | Energy cost Stabilized - Non-stabilized ~ Arm swing, 1 \| Participant | -0.02 | 0.55 | - | - | -0.18 | **0.01** | 3 | 4 | 134.59 | 150.08 | -63.30 | 1 vs 3 | 3.80 | 0.05 |

Our results showed that the model in which both Step width and Arm swing were included, did not provide significant variations compared to the models in which Step width or Arm swing were included separately (**Table S9**). The combination of Step width, Arm swing, and Rope length provided a correlation of −0.85 between Arm swing and Rope length. To avoid invalid results due to collinearity, we did not report the results of this regression analysis.

## Footnotes

1 This study was excluded from our meta-regression analysis because the individual participant data of this study were not accessible.

